# ‘Mini analysis’ is an unreliable reporter of synaptic changes

**DOI:** 10.1101/2024.10.26.620084

**Authors:** Ingo H. Greger, Jake F. Watson

## Abstract

Analysis of miniature postsynaptic currents (mPSCs, or ‘minis’) is one of the most extensively employed approaches to determine the functional properties of synaptic connections. The popularity of this technique stems from its simplicity. Patch-clamp recording is used to observe spontaneous transmission occurring at synapses throughout a neuron’s dendritic tree. These events are analysed in an attempt to determine quantal synaptic parameters and changes during experimental manipulation. For decades, changes in the amplitude or frequency of mPSCs have been interpreted as evidence for specific synaptic modifications, including changes in presynaptic release sites, postsynaptic receptor abundance or even nanoscale changes in the alignment of synaptic machinery. However, at the majority of brain synapses, these events are small, with many undetectable due to recording noise. Here, using both simulated and experimental data, we demonstrate that interpreting synaptic changes from mPSC datasets is fundamentally fallible. Due to incomplete detection of event distributions, seemingly specific changes in mPSC amplitude or frequency falsely report actual synaptic changes. In addition, probabilistic detection of small events gives false confidence in the completeness of detection and inaccurate determination of quantal size. We not only demonstrate the dense pitfalls of mini analysis, but also establish a method for experimental detection of the detection limit, allowing more robust data analysis and scientific interpretation.

## Introduction

The brain is an immensely complex organ. Its cells are connected by trillions of synapses, each with unique properties, determined by countless reactions and protein interactions (Raghavachari and Lisman, 2004; Yasuda, 2017). As these connections are potential sites for information storage in the brain, decades of research has been performed to determine the functional properties and molecular mechanisms of synaptic transmission (Nicoll, 2017).

However, determining and quantifying the properties of individual synaptic connections is surprisingly difficult. Since first observations of ‘quantal synaptic properties’ (del Castillo and Katz, 1954), the biophysical parameters of transmission have been studied intensively. One extensively employed method for this involves recording ‘miniature postsynaptic currents’ (‘minis’/mPSCs: excitatory – mEPSCs, inhibitory – mIPSCs) or spontaneous postsynaptic currents (sPSCs), using whole-cell patch clamp electrophysiology (Isaacson and Walmsley, 1995; Malgaroli and Tsien, 1992; Zhang and Trussell, 1994). Spontaneous events (sPSCs) are the result of action potential-dependent neurotransmitter release from any cell connected to the recorded neuron. Miniature events, recorded during action potential blockade (typically through tetrodotoxin (TTX) application), are the result of spontaneous vesicle fusion at any synapse on the recorded neuron (Kavalali, 2015). Both events occur anywhere across a target neuron’s dendritic tree, and allow ensemble sampling of synaptic input properties for a cell of interest. Given the ease of patch-clamp recording from individual neurons, m/sPSC analysis is routinely employed by labs worldwide to determine synaptic properties or synapse-level effects of experimental manipulations (e.g. gene knockout (Matt et al., 2018), synaptic protein manipulation (Gutierrez-Castellanos et al., 2017; Watson et al., 2017), or neuromodulatory action (Choy et al., 2018)).

Multiple parameters will determine the distribution of recorded synaptic events (Jonas and Spruston, 1994). It is often incorrectly assumed that changes in mPSC frequency reflect purely presynaptic changes (i.e. changes in number of release events), while mPSC amplitude differences are due to postsynaptic alterations (i.e. changes in response strength), and is still quoted as such in papers to date. This oversimplification will almost certainly drive incorrect biological conclusions. Multiple pre- and postsynaptic factors can influence both the amplitude and frequency of mPSCs. However, the pitfalls with mPSC interpretation only begin here. Not only can the biological interpretation of mPSC data be easily mistaken, but empirical interpretation of event distribution changes can be similarly misunderstood.

At the majority of brain synapses, synaptic strengths (and therefore mPSC amplitudes) are small (0-50 pA) (Edwards, 1991). They (mostly) approximate a right-skewed lognormal distribution (Buzsáki and Mizuseki, 2014; Sahara and Takahashi, 2001; Zhang and Trussell, 1994), while patch clamp noise levels are comparatively large (typically 2–10 pA max to min values/ 1–5 pA standard deviation when low pass filtered to 10 kHz). This results in many synaptic events lying beneath the noise (Malgaroli and Tsien, 1992; Isaac et al., 1996; Mennerick and Zorumski, 1995; Wang et al., 2024). Here, using *in silico* mPSC simulations, we demonstrate that calculating and considering the detection limit for mPSC events is essential for correct interpretation of these recordings. First, we demonstrate that events below the detection limit are probabilistically detected dependent on their amplitude, giving rise to a ‘false’ distribution that misrepresents quantal size. Second, we show that average event amplitude and frequency are not discrete parameters. Changes in event amplitude are predominantly represented as a selective change in mPSC frequency. Finally, we demonstrate a method for estimating the detection limit, allowing more reliable interpretation of recorded data. Together, this study demonstrates the major risk of mPSC misinterpretation, facilitating more accurate future investigation and re-examination of historic datasets.

## Results

In order to determine the effects of the detection limit on recorded mPSC distributions, we established a pipeline for simulating mPSC recordings with realistic patch-clamp noise. Using simulated data allows full control of both mPSC properties and recording noise levels. We first sought to determine the effect of event amplitude on detection around the detection limit. mPSC events were simulated with varying rise time constants, decay time constants, and peak amplitudes following a lognormal distribution, approximating real-world data (**Figure 1**) (Bekkers and Stevens, 1989; Zhang and Trussell, 1994; Sahara and Takahashi, 2001; Pernía-Andrade et al., 2012). These events were randomly positioned across a simulated recording (1 Hz event rate) and embedded in coloured noise (**Figure 1A-B**). We next detected events in these simulated recordings using a standard, template-fit algorithm (Clements and Bekkers, 1997; Jonas et al., 1993). The amplitude of detected events also followed a lognormal-like distribution, however a large fraction of events were not detected (**Figure 1C**). Unsurprisingly, undetected events were of small amplitude, hidden in recording noise. As logically expected due to loss of small events, the mean mPSC amplitude was overestimated by detection through recording noise (**Figure 1D**, actual mean event amplitude: 7.6 ± 0.1 pA; detected mean amplitude: 10.0 ± 0.1 pA), while mPSC frequencies were underestimated (**Figure 1D**, actual frequency: 1 Hz; detected frequency: 0.64 Hz). By calculating which simulated events were missed, we determined the association between mPSC amplitude and its likelihood of detection (**Figure 1E**). Event detection falls away probabilistically with decreasing amplitude, following a sigmoidal relationship. This probabilistic detection of small events shapes the rising phase of the detected event histogram (**Figure 1C**). Despite missing over a third of events, the resulting detected events had a lognormal-like distribution, reminiscent of a complete distribution. To demonstrate that this curve shape is independent of the input distribution, we detected events from a uniform distribution of simulated mPSCs (**Supplementary Figure 1**), which also demonstrated a gradual loss of mPSCs as their amplitude approached 0 pA. Due to the overabundance of small events, errors in measured event amplitude and frequency were much stronger for a lognormal than a uniform input distribution (**Figure 1, Supplementary Figure 1**). Therefore, the nature of synaptic properties unfortunately lends itself to more noise affected analysis. With complete detection, the mode (peak) of mPSC distributions would represent synaptic quantal size (Sahara and Takahashi, 2001; Gordleeva et al., 2023). However, in our example, this value was overestimated by at least factor of two (actual mode: 2.65 pA, detected mode: 5.92 pA). Therefore, incomplete detection produces mPSC datasets that do not accurately represent the underlying physiology, and give false confidence in their biological relevance.

**Figure 1.**
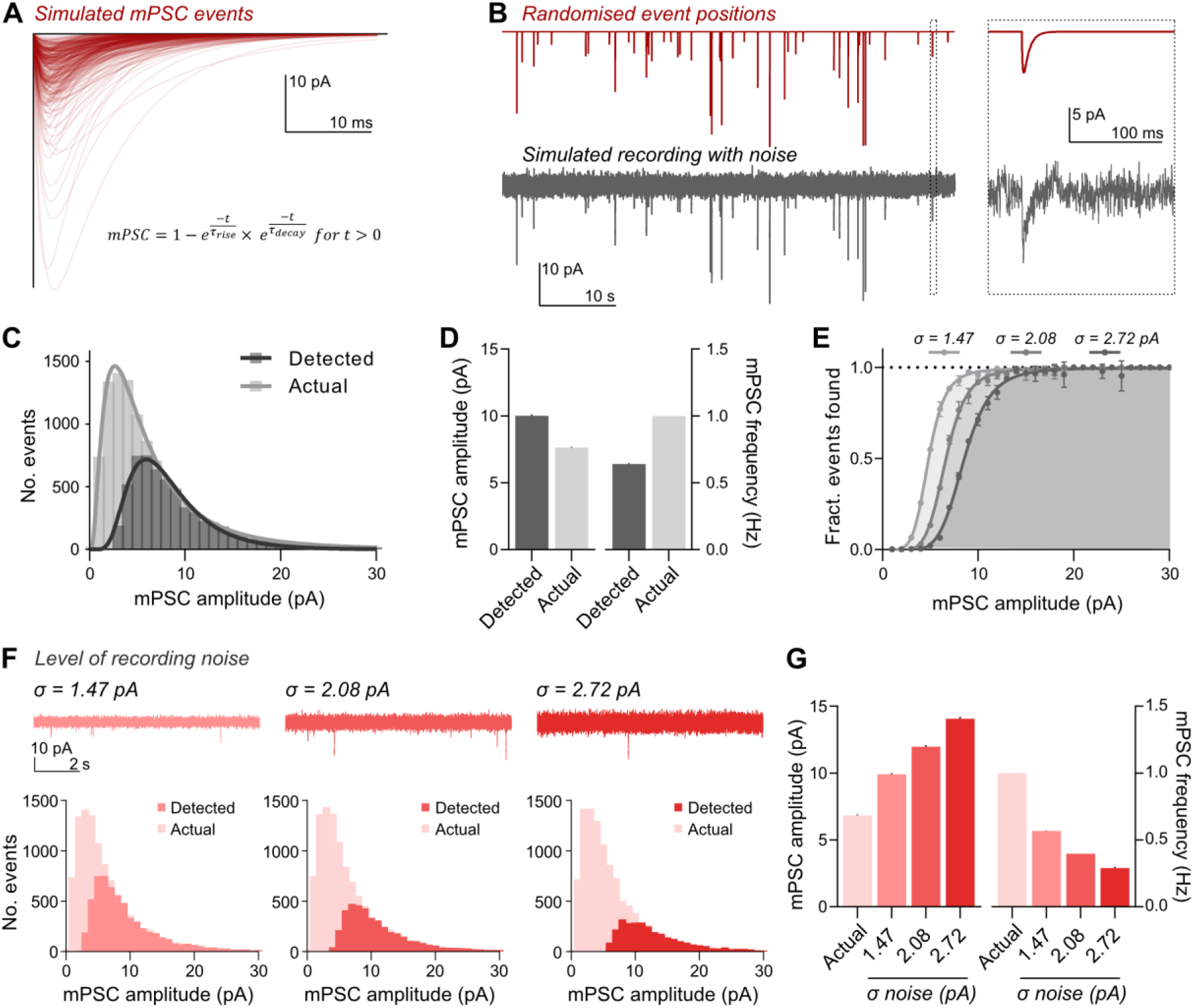
The detection limit creates a misrepresented event distribution. **A**, Simulated mPSC events (red) were created from a biexponential function (detailed), with randomised, realistic τrise, τdecay, and peak amplitudes. A sample of 300 events is depicted. **B**, Events were randomly distributed across a ‘time’ axis with controlled frequency (upper, 1 Hz) and embedded in simulated recording noise (lower). An expanded view (right) depicts a single event from the wider recording (boxed). **C**, Frequency distributions of simulated events before noise embedding (actual) and those extracted by template fit event detection (detected) demonstrate loss of small events after detection. **D**, Detected mean mPSC amplitude was overestimated (mean ± SEM; Actual: 7.6 ± 0.1 pA; Detected: 10.0 ± 0.1 pA), while event frequency was underestimated (Actual: 1 Hz; Detected: 0.64 Hz; n = 3 simulated recordings, 1 Hz events, 1 hr duration). **E**, Event detection follows a sigmoidal relationship, with a probabilistic decrease in event detection for smaller event amplitudes. Three levels of recording noise are displayed (σ = standard deviation of recording noise). **F**, The abundance of false negative introduction was dependent on recording noise. Increasing standard deviation of noise (example traces, upper) decreased the proportion of detected events (lower). **G**, Increased recording noise increased the mean detected event amplitude (mean; Actual, 6.8 ± 0.1 pA; σ = 1.47, 9.9 pA ± 0.1; σ = 2.08, 12.0 ± 0.1 pA; σ = 2.72, 14.1 ± 0.2 pA), while decreasing frequency (Actual, 1 Hz; σ = 1.47, 0.57 Hz; σ = 2.08, 0.40 Hz, σ = 2.72, 0.29 Hz).

We next modulated the detection limit by increasing the standard deviation of simulated noise. This caused a curve shift in the distribution of detected mPSCs (**Figure 1E-F**), and increased the misrepresentation of both mean mPSC amplitude and frequency (**Figure 1G**, see also **Supplementary Figure 1**). Strikingly, just a 0.6 pA increase in the standard deviation of recording noise caused a 21 % increase in the measured mean amplitude (9.9 to 12.0 pA mean amplitude), and an 30 % decrease in recorded frequency (0.57 to 0.40 Hz frequency) (**Figure 1G**). The level of noise therefore has a dramatic effect on the observed synaptic parameters. In real world recordings, noise is not just a result of recording setup. Noise levels vary depending the cell type, cell state, or even within and between individual recordings due to cell health, seal integrity and recording quality. Therefore, when comparing mPSC data between experimental groups, it is essential to compare recording noise of all cells across datasets.

### Detecting changes in synaptic properties

We next sought to determine how *changes* in mPSC properties are observed through the lens of recording and detection. Theoretically, a pure effect on event amplitude would produce a shift or scaling of the amplitude distribution along the *x*-axis, while frequency changes would induce a scaling of the distribution in *y* (**Figure 2A**). Therefore, in theory, visualising recorded event distributions should allow simple understanding of underlying synaptic changes. Indeed this is certainly the case when all events are detected, for example large events which occur well beyond the detection limit (e.g. action potential-dependent EPSCs at cochlear nuclei or Calyx of Held, (Sahara and Takahashi, 2001; Zhang and Trussell, 1994)). However, at the majority of synapses, and for almost all single-vesicle induced mPSCs, this is not the case.

**Figure 2.**
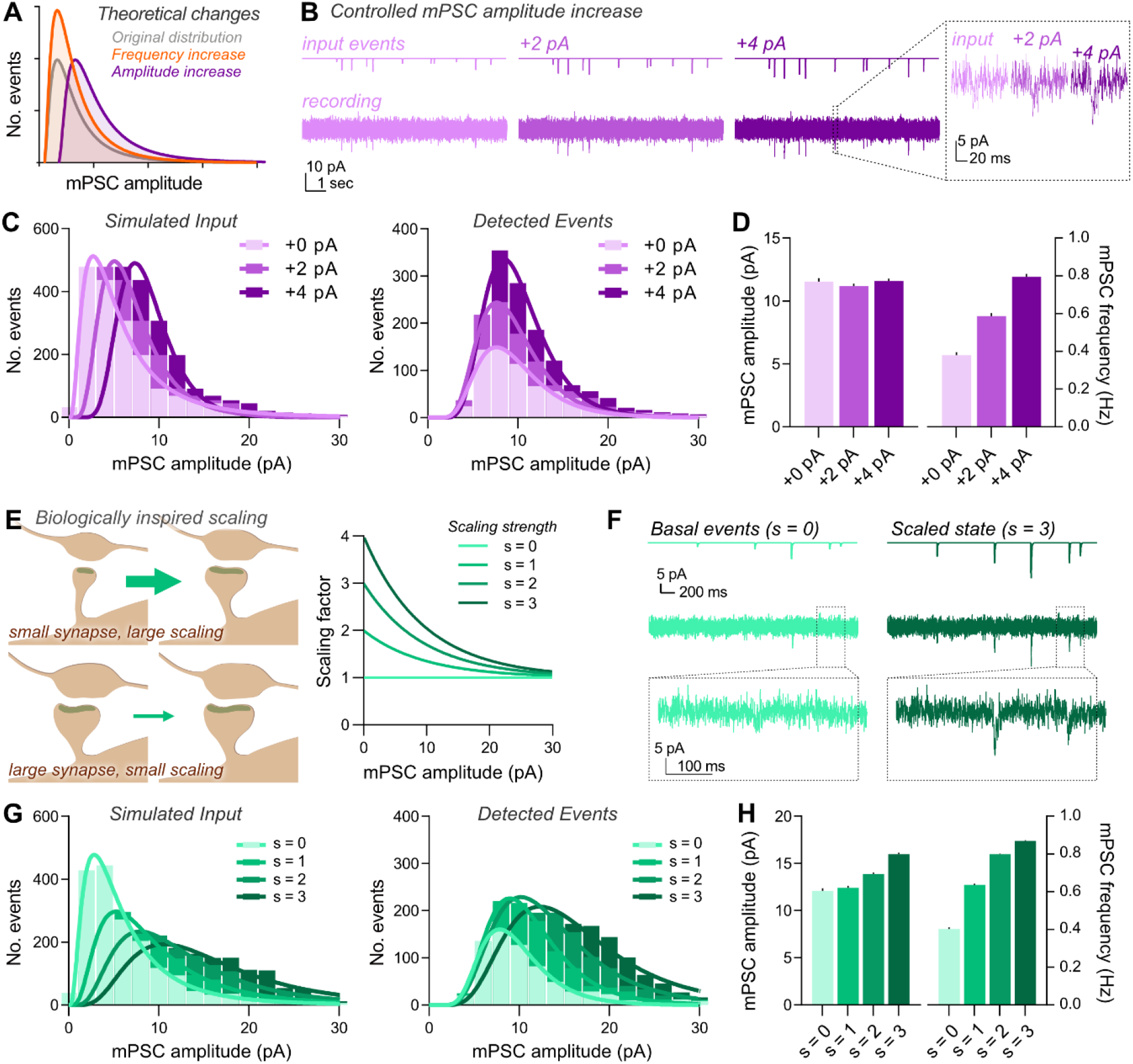
Incomplete event detection results in misidentification of mPSC changes. **A**, Theoretical changes in mPSC amplitude distributions for pure changes in frequency (orange) or amplitude (purple). **B**, Example traces of simulated mPSC recordings, with increasing event amplitudes (+2 pA or +4 pA). **C**, Simulated input events demonstrated an expected *x*-axis shift by increasing all event amplitudes, while distributions of detected events did not display *x*-axis shifting, despite pure amplitude modification. **D**, Mean detected event amplitudes were not altered when increasing the amplitude of input events (mean ± SEM; ‘+0 pA’, 11.5 ± 0.3 pA; ‘+2 pA’, 11.2 ± 0.2 pA; ‘+4 pA’, 11.6 ± 0.2 pA), while large changes in frequency were observed (right; ‘+0 pA’, 0.38 Hz; ‘+2 pA’, 0.59 Hz; ‘+4 pA’, 0.80 Hz). **E**, Principle (left), and model (right) of biologically inspired scaling rule, where small synapses were more strongly increased than larger synapses. Tested scaling factors (s) are depicted, where s = 0 corresponds to the unmodified distribution. **F**, Representative event simulations, depicting embedded events (upper), and simulated recording (lower) demonstrating basal (s = 0) and scaled recordings (s = 3). **G**, Input event distributions before (s = 0) and after applying scaling rules (left). Detected events from simulated recordings showed an increase in the distribution peak at low scaling (s = 1), before x-axis shifts were also observed with strong scaling (s = 3). **H**, Mean detected mPSC amplitude only increased for strong scaling factors, while observed mPSC frequency increased across all datasets (amplitude: s = 0, 12.1 ± 0.3 pA; s = 1, 12.4 ± 0.2 pA; s = 2, 13.9 ± 0.2 pA; s = 3, 16.0 ± 0.2 pA; frequency: s = 0, 0.4 Hz; s = 1, 0.64 Hz; s = 2, 0.80; s = 3, 0.87).

We repeated our mPSC simulations, but modified the event template to increase the peak amplitude of every event by either 2 pA or 4 pA before noise embedding (**Figure 2B**). This manipulation simulates a specific scaling in mPSC amplitude, independent of original synaptic weight. As expected, distribution histograms for the simulated events shift in *x* (**Figure 2C**). Histograms of *detected* events however, do not. We instead observe a stretching of the event distribution in the *y*-axis, which is reminiscent of a pure increase in mPSC frequency (**Figure 2C**). Despite simulating a pure increase in mPSC amplitude, the frequency of detected events increased, while mean amplitude was comparable (**Figure 2D**). Due to incomplete detection, small events emerge from the noise to both increase the detected frequency, and counteract the amplitude increase of previously detected events. Our data demonstrate that mPSC frequency and amplitude are not independent variables when working close to the detection limit. As a result, specific effects on either mPSC frequency or amplitude are unreliably reported, and definitive conclusions can only be made from datasets with complete event detection or through detailed analysis of recorded distributions.

We sought to corroborate these findings using a more biologically realistic scaling model. Events were scaled by a factor inversely proportional to their initial amplitude, simulating saturating potentiation and maximal effects on small synapses, as observed in biological systems (Kaneko et al., 2011) (**Figure 2E-F**). With weaker scaling, specific changes to detected event frequency occurred (**Figure 2G-H**). Only with more robust scaling was a shift in the detected event distribution towards higher amplitudes observed (**Figure 2G**), yet this was accompanied by a >2-fold increase in mPSC frequency (**Figure 2H**). These results further confirm the interdependence of mPSC frequency and amplitude for noise-embedded events, and the strong sensitivity of mPSC frequency measurements to underlying changes in amplitude.

### Detecting changes in event frequency

To complete this analysis, we applied controlled changes in mPSC frequency (**Figure 3**). We simulated mPSC recordings with frequencies of 1, 2 and 3 Hz. Despite the limits of detection detailed previously, both event distributions and mean frequencies reflected these changes, albeit strongly underestimating true frequencies (**Figure 3D**). Mean mPSC amplitudes showed little change. While this data confirms that pure mPSC frequency changes can be reliably followed, such effects cannot be concluded from ‘real-world’ data, as observed changes in mPSC frequency could occur through biological changes in either amplitude or frequency.

**Figure 3.**
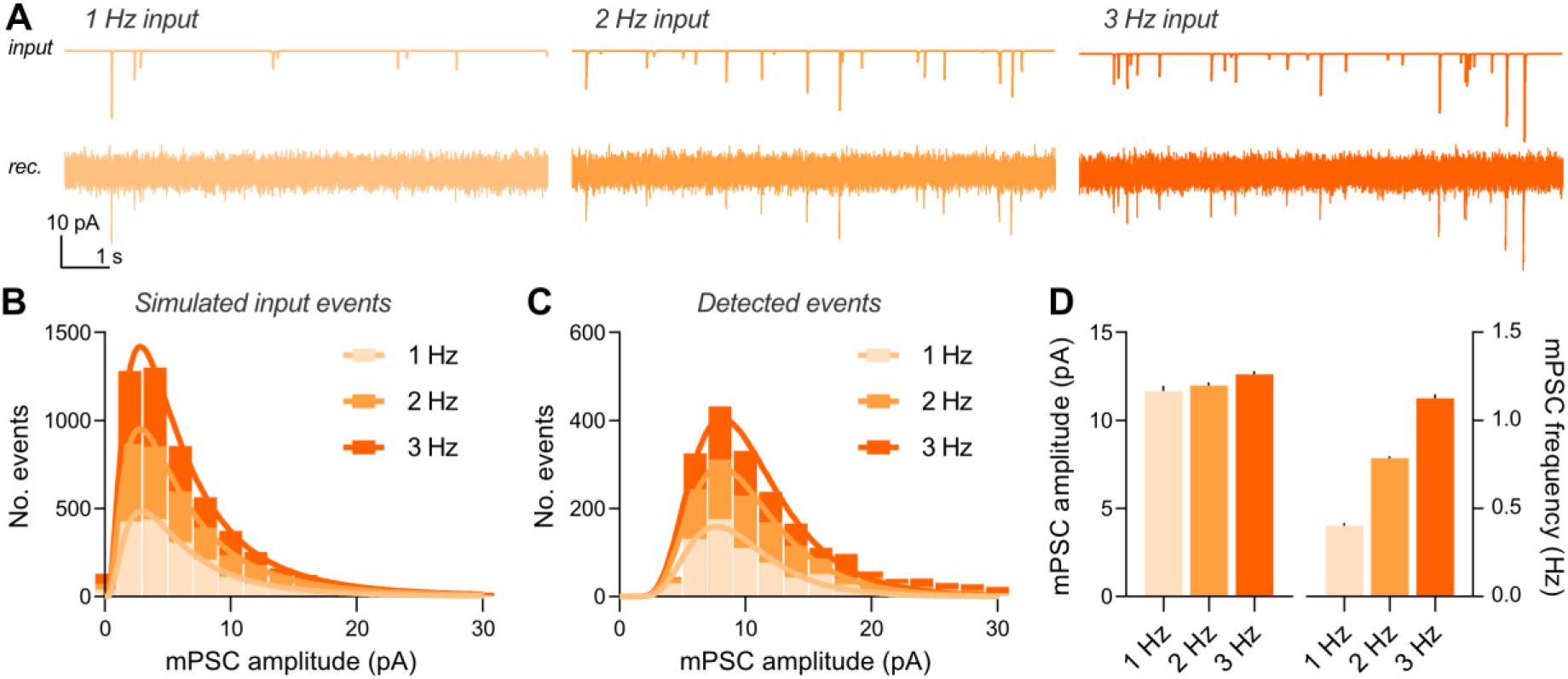
Measurement of mPSC frequency changes. **A**, Example traces from simulated input at increasing mPSC frequency (1-3Hz). **B**, Input event histograms skewed purely in the y-axis. **C**, Detection of mPSC frequency changes were not distorted by detection. **D**, Input frequency change produced specific changes in detected frequency (right; 1 Hz, 0.4 Hz; 2 Hz, 0.8 Hz; 3 Hz, 1.1 Hz), and not mean amplitude (left; 1 Hz, 11.7 ± 0.3 pA; 2 Hz, 12.0 ± 0.2 pA; 3 Hz, 12.6 ± 0.2 pA).

### Real-world mPSC data contains hidden distributions

While simulated data are valuable for testing mPSC analysis in a controlled system, we sought to assess the unreliability of mPSC detection with experimentally recorded mPSCs. Many pharmacological manipulations are known to enhance synaptic transmission, but due to the complexity of synaptic transmission, it is difficult to confidently change transmission amplitude without frequency effects. However, this can be achieved electronically. Recording mPSC datasets from the same neuron at two holding potentials allows specific and calculable modification of mPSC amplitude, without influencing mPSC frequency, by increasing the driving force for ion flow across the membrane. This approach has been previously employed as such for an experimental control (Malgaroli and Tsien, 1992; Manabe et al., 1992), and to dissect synaptic parameters (Chen et al., 2015). We performed mEPSC recordings from CA1 pyramidal neurons in mouse organotypic slice cultures at both -60 mV and -80 mV holding potentials (in the presence of 1 μM tetrodotoxin to block action potential generation, 10 μM SR-95531 to isolate mEPSCs, and 100 μM D-APV to isolate the AMPAR current) (**Figure 4A**). Given the variability of real world synaptic data, we sampled equal length recordings of mEPSCs at both holding potentials from every included cell, and proceeded with mEPSC detection and analysis. The difference in holding potential should cause a specific 1.33-fold scaling in mEPSC amplitude. However, we observed no change in mean mEPSC amplitude and instead observed an increase of mEPSC frequency (**Figure 4C**). Event distributions show y-axis scaling as expected from a frequency change, directly replicating our theoretical analysis (**Figure 4B**). Recorded cells showed no difference in noise levels between conditions, ensuring our conclusions were not affected by detection variability (**Figure 4D**). These observations corroborate our simulations, demonstrating that real-world mPSC analysis is confounded by the detection limit, and that a significant portion of mPSC events occur below the detection limit at hippocampal CA1 synapses.

**Figure 4.**
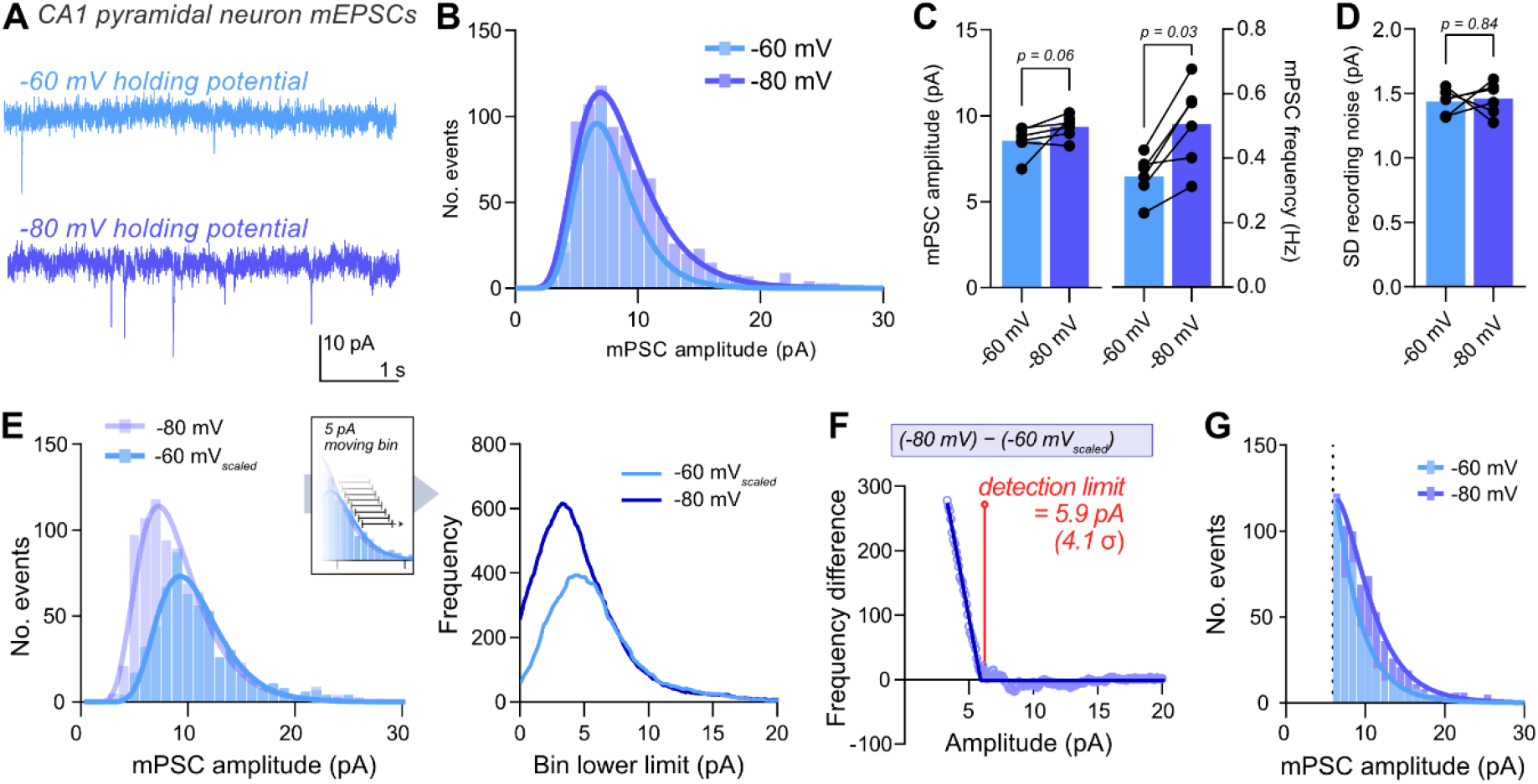
Demonstrating the fallibility of real-world mPSC data, and detection of the detection limit. A, Example mEPSC traces recorded from CA1 pyramidal neurons at -60 and -80 mV holding potentials. **B**, Histograms of recorded events demonstrated an overall frequency increase (lognormal fit, 2 pA bin width). **C**, Observed mean event amplitudes were not affected by holding potential driven amplitude increase (−60 mV, 8.44 ± 0.17 pA; -80 mV, 9.40 ± 0.16 pA; n=6 cells; Wilcoxon matched-pairs signed rank test, p = 0.06), yet observed frequency was increased (−60 mV, 0.34 ± 0.03 Hz; -80 mV, 0.51 ± 0.05 Hz; n = 6 cells; Wilcoxon matched-pairs signed rank test, p=0.03). **D**, There was no difference in the standard deviation (SD) of recording noise between conditions (−60 mV, 1.44 ± 0.04 pA; -80 mV, 1.46 ± 0.08 pA; n=6 cells; Wilcoxon matched-pairs signed rank test, p=0.84). **E**, Scaling the -60 mV dataset by 1.33 produced overlapping distributions at high amplitudes. These datasets were resampled by a 5 pA-sliding bin at a resolution of 0.1 pA (right). **F**, Plotting the difference between resampled datasets allowed determination of the amplitude at which events begin to be ‘lost’ at -60 mV but not at -80 mV. The frequency difference curve was fitted with a broken stick relationship, which approximated the detection limit (5.9 pA or 4.1 σ of recording noise). **G**, Distribution of recorded mEPSCs after application of the detection limit as a cut-off. Histogram bins start from the detection limit. No peak to event distribution was observed in the recorded range.

### Experimental detection of the detection limit

We have demonstrated the dramatic fallibility of mPSC analysis for understanding and interpreting biological effects. Using holding potential scaling, we were able to ‘visualise’ mPSC events that exist, but were hidden beneath the detection limit when recording at -60 mV. We next sought to determine a means to improve the reliability of mPSC analysis, through determination of the event detection limit. mEPSCs recorded and detected using a -60 mV holding potential were scaled to their expected amplitude at -80 mV (x1.33), and plotted alongside our ‘ground truth’ -80 mV mEPSC dataset (**Figure 4E**). While these distributions overlay almost perfectly at high event amplitudes (away from the detection limit), the -60 mV dataset showed lower frequency of small amplitude events. Our -80 mV dataset was also likely be incomplete at small amplitudes, yet contains a more complete representation than that recorded at -60 mV. Therefore, the point at which these two curves diverge is the amplitude at which mPSCs begin to become undetected, i.e. the detection limit. A sliding bin histogram of mPSC events was plotted for -80 mV, and scaled -60 mV datasets (**Figure 4E**). Calculating the difference between these curves (subtraction of -60 mV_scaled_ from -80 mV) produces a biphasic curve, where high amplitudes can be fit with a y=0 curve (no difference in event detection), yet low amplitudes approximate a linear relationship with negative gradient (**Figure 4F**). This graph can be approximated with a ‘broken stick’ curve, where the break point represents the lowest amplitude at which zero false negatives are recorded, or the ‘detection limit’ (**Figure 4F**). In our dataset, the detection limit was estimated at 5.9 pA, or 4.1 times the standard deviation of recording noise (1.43 pA, σ denotes standard deviation).

Finally, we used this knowledge of the detection limit to re-examine recorded event distributions. We applied the recorded detection limit as a cut off for included events, and replotted recorded datasets with the minimum bin value beginning at this cut-off (**Figure 4G**). We were unable to fit a peak to our real-world mEPSC dataset, demonstrating that the modal event remains below the detection limit. Estimates of quantal size or in depth interpretation of distribution changes would not be possible from such data.

## Discussion

The properties of chemical synapses are highly diverse, both across the brain and at the level of individual neurons. For this reason, understanding the changes in synaptic physiology underlying brain function requires robust methods for their study. Analysis of mPSCs has the potential to provide information about heterogeneous synaptic efficacies across the neuronal dendritic tree. This approach is powerful and technically simple, but the pitfalls of data analysis and interpretation are deep, hidden, and not widely appreciated. We have demonstrated these issues using both simulated and experimental datasets, suggesting analyses for careful interpretation of recorded data. While this manuscript focuses on mini analysis (mPSCs), these concepts are directly applicable to sPSC recordings, or analysis of any detected event that has close proximity to noise levels.

### Empirical interpretation of mPSC datasets

It would be logical to assume that measured changes in mean event frequency represent underlying changes in mean event frequency, and similarly event amplitudes. However, our data demonstrates that this is a false assumption. When events are embedded in recording noise, changes in mPSC amplitude are more robustly detected as changes in frequency than mean event amplitude (**Figures 2 & 4**). Therefore, even specific observed changes in mPSC frequency could be caused by underlying changes in either frequency or amplitude. This interdependence was well appreciated in the early years of mini analysis (Diamond and Jahr, 1995; Manabe et al., 1992; Mennerick and Zorumski, 1995; Yamada et al., 1993), yet now appear widely forgotten. Not only are changes in mPSC frequency an unreliable indicator of biological changes, but changes in mean amplitude are more sensitive to the level of recording noise than to actual underlying synaptic changes. Without careful analysis of recorded noise levels between conditions, these effects have the potential to cause regular misinterpretation of synaptic changes from mini analysis.

While our results demonstrate the importance of interpreting event distributions rather than averages values, distributions can be similarly misleading. If complete distributions are seen above the noise level, interpreting mPSC changes is not an issue, and even mean values will reliably report underlying biological changes. For the majority of brain synapses, this is not the case, including the most intensively studied excitatory connections across the hippocampus and cortex. We show here that the modal mEPSC of CA1 pyramidal neurons lies beneath recording noise in our hands (<5.9 pA).

Unfortunately, the detection limit is not simply a sharp cut-off. Small events are lost with increasing likelihood the smaller they are, creating a false ‘peak’ to event amplitude distributions. The resulting distributions strongly resemble an expected lognormal-like profile of synaptic events, giving false confidence that measured data fully represent the underlying biology. Quantal analysis from mPSC data is also therefore problematic, unless either clear evidence of quantal properties are observed e.g. (Paulsen and Heggelund, 1994), or the peak of the event distribution can be unequivocally observed above the detection limit. Even then, the level of variability observable at neuronal synapses will often preclude simple interpretation of such results (Clements, 1991; Malinow, 1991).

To facilitate data analysis, we present a means to estimate the detection limit. Using different voltage clamp holding potentials, electronically scaled mPSCs can be compared to determine the amplitude at which events reliably emerge from noise (**Figure 4**). By using this approach, the amplitude at which mPSC analysis becomes unreliable can be determined for a certain recording setup/system. Event distributions can then be attenuated so as to analyse only those falling above this point, preventing false peaks in amplitude distributions. Our detection limit estimation from real world data (4.1σ) is directly in line with previous estimates. Clements and Bekkers predicted that 4 × σ would eliminate false negative detection (Clements and Bekkers, 1997). Therefore, where empirical detection limit measurement is not possible, 4σ may be an appropriate cut off for mPSC analysis. It is important to note that when binning data attenuated at a calculated detection limit (e.g. histogram presentation), lower bin limits must start precisely at the event cut-off, therefore most likely requiring non-integer edge values.

Recording noise is dependent not just on setups, but also individual cells. The quality of sealing and membrane integrity during whole-cell recordings will influence the standard deviation of noise, and event detection will be dependent in turn on these parameters. It is therefore important that analysed mPSC recordings not only have as low noise as possible, but also that experimental groups have equivalent noise levels between cells, and that lower quality recordings are discarded. When experimental groups have different levels of noise, analysis can be reliably performed by applying the detection threshold from the highest noise dataset to all conditions.

### Biological interpretation of mPSC changes

Interpretation of ‘mini data’ extends further than only numbers and distributions. As we understand more about the plasticity of synapses, it becomes clear that the historic doctrine of ‘mPSC amplitude changes = postsynaptic’ and ‘mPSC frequency changes = presynaptic’ is a gross oversimplification. Multiple factors of both pre and postsynapse could biologically alter mPSC frequency and amplitude (**Figure 5**). Changes in mPSC amplitude could be caused postsynaptically, by changes in neurotransmitter receptor abundance or conductance (Kessels and Malinow, 2009; Malenka and Nicoll, 1999), but also presynaptically or transsynaptically, through vesicle properties or alignment (Scheefhals and MacGillavry, 2018). Similarly, changes in the number of active release sites could change mPSC frequency (Malgaroli and Tsien, 1992), but postsynaptic unsilencing (Isaac et al., 1996) could also enact a postsynaptic change in event frequency. Neuronal properties such as dendritic event location and cable properties will also influence mPSC properties. Each of these factors can add significant complexity to the distribution of synaptic events recorded from across a neuron’s dendritic tree. Therefore, interpreting specific effects, such as changes in synaptic nanoarchitecture, would need both low noise recordings and particularly careful data analysis.

**Figure 5.**
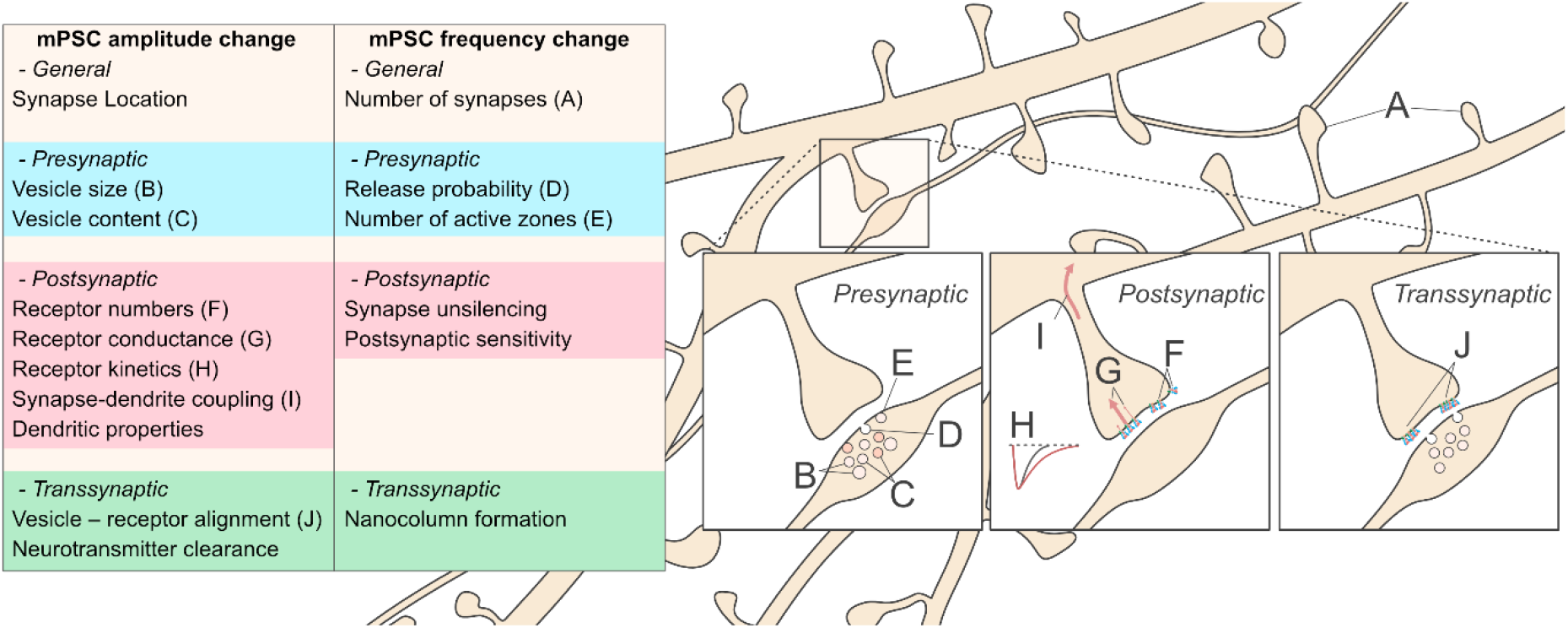
Overview of possible factors influencing mPSC changes. Both mPSC frequency and amplitude may be altered by a range of pre (blue), post (red) and transsynaptic (green) changes, complicating interpretation of recorded observations.

‘Mini analysis’ has become widespread due to the ease of single-cell recording to acquire synaptic insights. While powerful, simple, and widely employed, this approach is prone to misinterpretation. We hope that this study can aid robust data analysis, strengthening insightful synaptic physiology and neuroscience research.

## Materials and Methods

### mPSC event simulation

Event simulation and recording noise generation was performed following (Pernía-Andrade et al., 2012). All simulated recordings were generated in MATLAB. mPSCs were simulated as biexponential functions formed of a rising exponential (τ_rise_) and decaying exponential (τ_decay_) following:

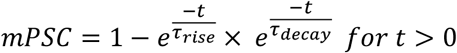

where *t* spanned 70 ms. τ_rise_ and τ_decay_ were randomly selected for each event from a lognormal distribution (Derkach et al., 1983) of realistic possible values (τ_rise_ μ: 0.2, σ: 0.25; τ_decay_ μ: 1.7, σ: 0.4), with τ_rise_ constrained between 0.3 < τ < 2.5 ms, and τ_decay_ between 1 < τ < 25 ms. Resulting curves were scaled to peak amplitudes randomly sampled from a lognormal distribution of realistic peak amplitudes (peak μ: 1.6, σ: 0.8). For scaled amplitude datasets, scaling factors (amplitude addition or multiplication) were applied to target peak amplitudes prior to curve scaling.

Biologically relevant synaptic scaling was implemented by multiplying individual peak amplitudes by a scaling factor dependent on the initial amplitude. This scaling factor followed an exponential decay relationship as follows:

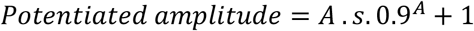

where *A* is the initial amplitude and *s* is the scaling factor of potentiation (varied between 0 and 3).

After their generation, events were assigned locations on the recording trace with uniform randomness, with the total number of assigned events determined from the product of desired frequency and recording length. This ‘noise free’ trace was then embedded in simulated recording noise. Events were simulated at 1 Hz unless otherwise stated. Generation of random locations or values was performed using MATLAB functions.

### Recording noise simulation

Simulated patch-clamp noise was produced by first, generating ‘white noise’ of variable standard deviation from normally distributed numbers centred on zero, before inclusion of a 1/f ‘pink noise’ component and Gaussian filtering the resulting mixed noise at 1000 Hz (**Supplementary Figure 2**). The resulting noise has a comparable frequency spectrum to ‘real world’ patch clamp recording noise. The standard deviation of final noise was 2.08 pA unless otherwise stated.

### Event detection

Event detection employed a standard template search approach, based either in Clampfit, or MATLAB. MATLAB template detection employed the ‘minidet’ function from the Biosig toolbox (Schölgl et al., 2011), based on Jonas et al. 1993. Detected events were extracted from recordings or simulations and fit with a biexponential function consisting of a rising and decaying phase, and peak amplitude was taken as the minimum of the curve fit. This approach prevents error in peak measurement caused by recording noise, which is large for small events close to the noise level (e.g. mPSCs). The equation of fitted curved was:

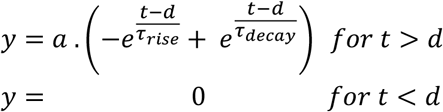

where *a* is the peak scaling factor, and *d* is the event onset time. mPSC frequency is calculated as the number of events detected per unit time (Hz). The fraction and properties of false negative (‘missed’) events was calculated by comparison of detected event times, with encoded event times.

The detection limit was estimated by sampling datasets of events recorded at either -60 mV (scaled by 1.33) and -80 mV holding potentials with moving bin of 5 pA width at resolution of 0.1 pA. Using the resulting frequency curves, the -60 mV_scaled_ dataset was subtracted from the -80 mV dataset and plotted against the bin lower limit. This graph was fit with a ‘broken stick’ curve, where:

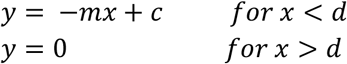

and ‘d’ represents the minimal amplitude at which zero false negatives are recorded, or the ‘detection limit’.

### Organotypic culture

Organotypic slice cultures were made using the Stoppini method (Stoppini et al., 1991), as described in (Watson et al., 2017). All procedures were carried out under PPL 70/8135 in accordance with UK Home Office regulations. Experiments conducted in the UK are licensed under the UK Animals (Scientific Procedures) Act of 1986 following local ethical approval.

Hippocampi from P6-8 C57/Bl6 mice of either sex were isolated in high-sucrose Gey’s balanced salt solution containing (in mM): 175 Sucrose, 50 NaCl, 2.5 KCl, 0.85 NaH_2_PO_4_, 0.66 KH_2_PO_4_, 2.7 NaHCO_3_, 0.28 MgSO_4_, 2 MgCl_2_, 0.5 CaCl_2_ and 25 glucose at pH 7.3. Hippocampi were cut into 300 μm thick slices using a McIlwain tissue chopper and cultured on Millicell cell culture inserts (Millipore Ltd) in equilibrated slice culture medium (37°C/5% CO2). Culture medium contained 78.5% Minimum Essential Medium (MEM), 15% heat-inactivated horse serum, 2% B27 supplement, 2.5% 1 M HEPES, 1.5% 0.2 M GlutaMax supplement, 0.5% 0.05 M ascorbic acid, with additional 1 mM CaCl_2_ and 1 mM MgSO_4_ (all from Thermo Fisher Scientific; Waltham, MA). Medium was refreshed every 3–4 days, and recordings were performed 10-12 days *in vitro*.

### Electrophysiology

Hippocampal slices were submerged in room temperature aCSF containing (in mM): 125 NaCl, 2.5 KCl, 1.25 NaH_2_PO_4_, 25 NaHCO_3_, 10 glucose, 1 sodium pyruvate, 4 CaCl_2_ and 4 MgCl_2_ at pH 7.3 and saturated with 95% O_2_/5% CO_2_. Excitatory mEPSCs were isolated by addition of 1 μM tetrodotoxin, 10 μM SR-95531 and 100 μM D-APV.

Borosilicate pipettes (3-6 MΩ tip resistance when filled with intracellular solution), were filled with intracellular solution containing (in mM): 135 CH_3_SO_3_H, 135 CsOH, 4 NaCl, 2 MgCl_2_, 10 HEPES, 4 Na_2_-ATP, 0.4 Na-GTP, 0.15 spermine, 0.6 EGTA, 0.1 CaCl_2_, at pH 7.25. CA1 pyramidal neurons were recorded in the whole-cell patch clamp configuration. Signals were acquired using a Multiclamp 700B amplifier (Axon Instruments) and digitized at 10 kHz using a Digidata 1440 A interface (Axon Instruments). Recordings during which the series resistance varied by more than 20% or exceeded 20 MΩ were discarded. Recordings were not corrected for the liquid junction potential.

### Data analysis and visualisation

All data were analysed in MATLAB (R2022), plotted in GraphPad Prism 9, and presented using Affinity Designer 2. Bars and errors in figures and legends present mean +/-standard error of the mean unless otherwise stated.

## Acknowledgements

The authors are very grateful to Prof. Helmut Kessels and Hinze Ho for initial discussions that led to this study, Dr. Andrew Penn for constructive feedback on the project, and Profs. Peter Jonas and Roger Nicoll for feedback on the manuscript. This work was supported by Biological Services teams at both the Laboratory of Molecular Biology and Ares facilities. Funding was provided by the Medical Research Council (MRC - MC_U105174197 to I.H.G.) and Horizon 2020 through Marie Skłodowska-Curie Actions (MSCA-IF 101026635 to J.F.W.).

## Data and code availability

Mini detection software from the BioSig project was used, with code available at https://biosig.sourceforge.net/. All additional data and code are available from the corresponding author upon request.

## Supplementary Information

**Supplementary Figure 1.**
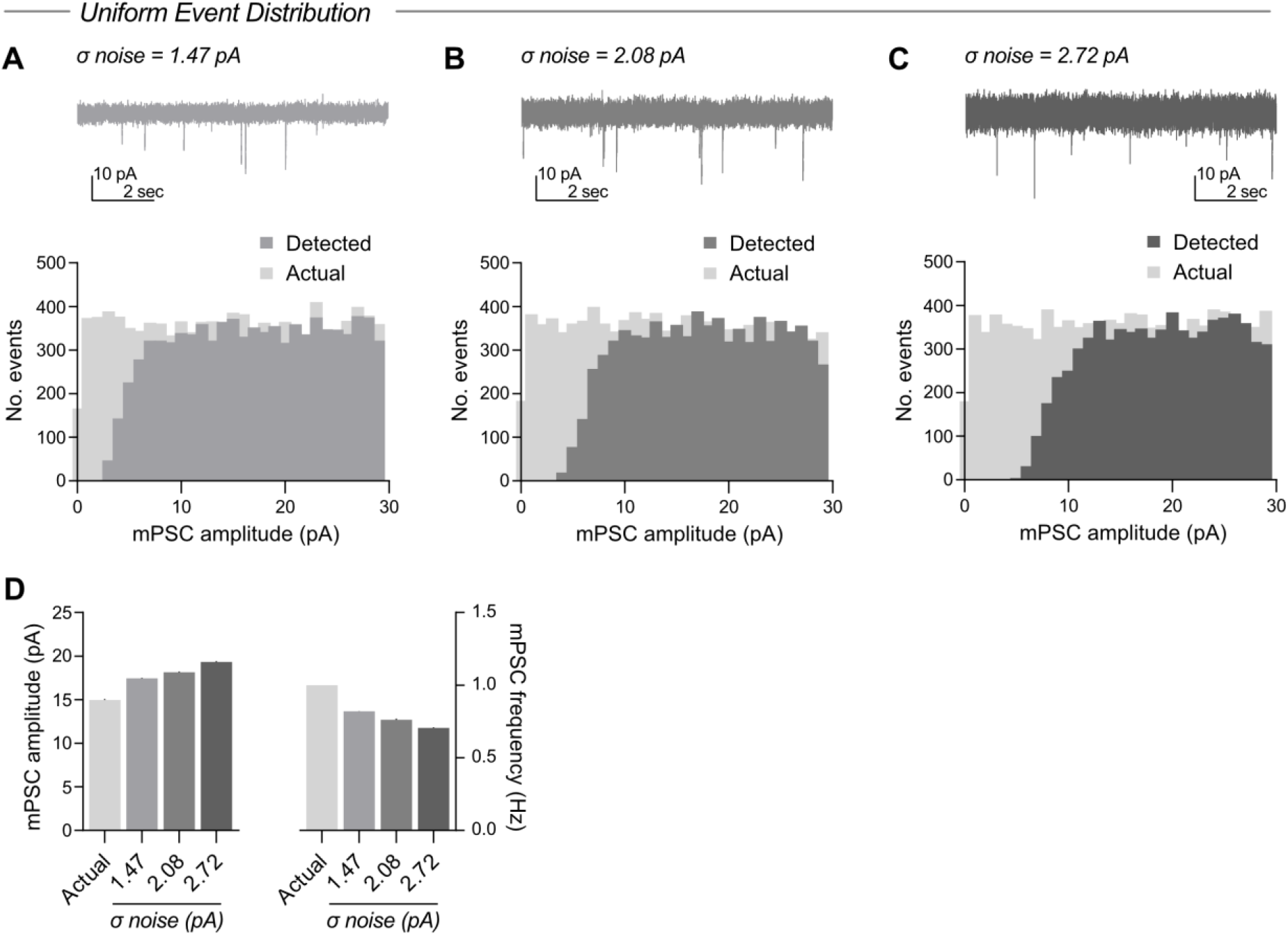
Event amplitude filtering by the detection limit. **A-C** Detection of events from a uniform distribution of simulated input amplitudes demonstrates probabilistic detection of small events, causing filtering of distributions by the detection limit. Increasing recording noise shifts this distribution to higher amplitudes. σ denotes standard deviation of recording noise. Mean event amplitudes and frequencies are presented in **D**.

**Supplementary Figure 2.**
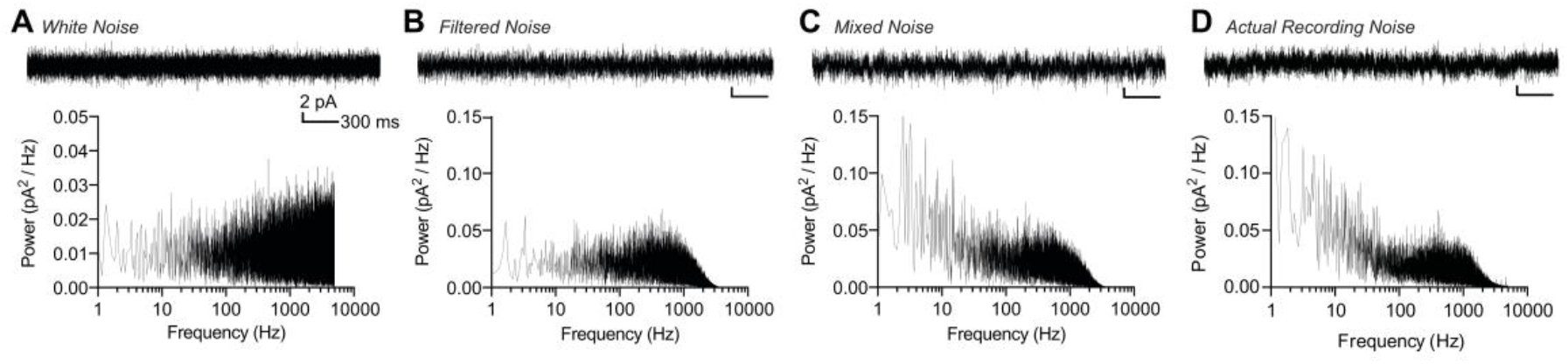
Frequency spectra for simulated recording noise. Example traces (upper) and frequency power spectra (lower) for unfiltered white noise (**A**), white noise filtered to 1000 Hz (**B**), mixed noise, containing both white and 1/f components filtered to 1000 Hz (**C**). **D**, Recording noise from experimental patch-clamp recording.

